# Expanding the spectrum of annexin A11 proteinopathy in frontotemporal lobar degeneration and motor neuron disease

**DOI:** 10.1101/2025.06.26.661831

**Authors:** Nikhil B. Ghayal, Richard J. Crook, Angita Jain, Gunveen Sachdeva, Shanu F. Roemer, Hiroaki Sekiya, Michael A. DeTure, Matthew Baker, Wouter De Coster, Bjorn E. Oskarsson, Keith A. Josephs, Rosa Rademakers, Marka van Blitterswijk, Dennis W. Dickson

## Abstract

Aggregation of TAR DNA-binding protein 43 (TDP-43) is strongly associated with frontotemporal lobar degeneration (FTLD-TDP), motor neuron disease (MND-TDP), and overlap disorders like FTLD-MND. Three major forms of motor neuron disease are recognized and include primary lateral sclerosis (PLS), amyotrophic lateral sclerosis (ALS), and progressive muscular atrophy (PMA). Annexin A11 (ANXA11) is understood to aggregate in amyotrophic lateral sclerosis (ALS-TDP) associated with pathogenic variants in *ANXA11*, as well as in FTLD-TDP type C. Given these observations and recent reports of *ANXA11* variants in patients with semantic variant frontotemporal dementia (svFTD) and FTD-MND presentations, we sought to characterize ANXA11 proteinopathy in an autopsy cohort of 379 cases with FTLD-TDP, as well as FTLD-MND and MND-TDP cases subclassified neuropathologically into PLS, ALS, and PMA. All FTLD-TDP type C cases had ANXA11 proteinopathy. However, ANXA11 proteinopathy was present in over 40% of FTLD-MND and in 38 out of 40 FTLD-PLS cases (95%), of which 80% had TDP type B or an unclassifiable TDP-43 proteinopathy and 15% had TDP type C. Genetic analyses excluded pathogenic *ANXA11* variants in all ANXA11-positive cases. We thus demonstrated novel forms of ANXA11 proteinopathy strongly associated with FTLD-PLS, but not with TDP type C or pathogenic *ANXA11* variants. Given the emerging relationship of ANXA11 in TDP-43 proteinopathies, we propose that TDP-43 and ANXA11 proteinopathy (TAP) comprises the molecular pathology of cases with abundant inclusions that are co-immunoreactive for both proteins and we subclassify three types of TAP based on distinct clinical and neuropathologic features.

## Introduction

Frontotemporal lobar degeneration (FTLD) and motor neuron disease (MND) are two overlapping neurodegenerative disorders characteristically associated with an accumulation of filamentous protein aggregates. FTLD encompasses many focal cortical and subcortical pathologies that often manifest clinically as frontotemporal dementia (FTD), primary progressive aphasia (PPA), and atypical parkinsonian disorders[30]. On the other hand, MND encompasses clinical and pathologic manifestations that variably affect upper motor neurons in the primary motor cortex, their descending corticospinal tracts, and lower motor neurons in the brainstem or spinal cord[22]. Several forms of MND are widely recognized, including primary lateral sclerosis (PLS) that predominantly affects upper motor neurons, amyotrophic lateral sclerosis (ALS) that affects upper and lower motor neurons, and progressive muscular atrophy (PMA) that predominantly affects lower motor neurons. Up to 50% of MND patients have cognitive impairment and motor neuron dysfunction is detected in close to 15% of FTD patients (FTD-MND). Overlapping pathologic features in many patients suggests that FTLD and MND comprise a spectrum of disorders with shared pathomechanisms[14, 54, 55, 77].

Nearly two decades ago, most tau-negative, ubiquitinated inclusions were found to contain aggregated TAR-DNA binding protein 43 (TDP-43) in patients with FTLD (FTLD-TDP), MND (MND-TDP), and FTLD-MND[70, 80]. Following this discovery, three common subtypes of TDP-43 proteinopathy with characteristic clinical, pathologic, and genetic features were recognized: TDP type A is associated with behavioral variant FTD (bvFTD), pathogenic variants in *granulin* (*GRN*), and the presence of neuronal cytoplasmic inclusions (NCI), short neurites, and neuronal intranuclear inclusions (NII); TDP type B is common in patients with clinical MND, FTD-MND, and carriers of pathogenic repeat expansions in *C9orf72* (*C9orf72*-RE), and is characteristically an NCI-predominant pathology; TDP type C often presents with semantic variant PPA (svPPA) and FTD (svFTD), lacks known genetic contributions, and is characterized pathologically by long, rope-like neurites[68]. Notably, we previously identified TDP type C in several patients who had FTLD with corticospinal tract and upper motor neuron-predominant degeneration consistent with PLS (FTLD-PLS)[16, 22, 34, 37]. In addition to TDP types A, B, and C, several less common types have been described: TDP type D associated with abundant NIIs and pathogenic variants in *VCP*, TDP type E associated with abundant grain-like inclusions and rapid clinical progression, and abundant perivascular and glial TDP-43 pathology associated with pathogenic *DCTN1* variants or Perry syndrome (TDP type P)[27, 45, 69, 91]. Many studies have reported unclassifiable TDP-43 proteinopathies with pathologic features of multiple TDP-43 types in patients who often have *C9orf72*-RE[3, 22, 31, 42, 56, 57, 68, 71, 86].

TDP-43 aggregation can also occur in the company of other aggregating proteins, such as tau, alpha-synuclein, or mutant proteins[26, 36, 38, 40, 41, 90, 94]. In such multi-proteinopathies, each aggregating protein typically accumulates in a distinct anatomical distribution in the brain (e.g., FTLD-TDP type A with primary age-related tauopathy). Despite this, similar anatomical distributions of inclusions containing TDP-43 and the molecular tether protein, annexin A11 (ANXA11), were recently described in MND patients harboring pathogenic variants in *ANXA11*[82]. Subsequent studies of *ANXA11* variant carriers have also described similar distributions of TDP-43 and ANXA11 inclusions in association with mainly MND-TDP, FTLD-MND, and inclusion body myopathy[33, 47, 67, 78, 79, 87]. Intriguingly, most studies have shown TDP-43 and ANXA11 labeling discrete, or separate, aggregates, and have less consistently shown TDP-43 and ANXA11 labeling the same aggregate.

Conversely, researchers initially investigating an *ANXA11* variant in an FTLD-TDP type C patient recently discovered that TDP-43 and ANXA11 consistently labeled the same aggregate in all TDP type C patients[78]. Remarkably, structural examination of TDP type C amyloid filaments by cryo-electron microscopy (cryo-EM) clarified this finding by revealing a heteromeric filament structure consisting of co-assembled TDP-43 and ANXA11 fragments[7]. In contrast, similar investigations in TDP types A and B have described homomeric filament structures consisting of distinct TDP-43 fragments[5, 6]. Consistent with these observations, ANXA11 inclusions were rare in TDP types A and B (<5%) in the former study[78]. Additionally, a separate study recently confirmed the presence of TDP-43 and ANXA11 proteinopathy in several TDP type C cases that had FTLD with upper motor neuron-predominant degeneration and corticospinal tract pathology[93].

Together, these reports suggest several roles for ANXA11 proteinopathy throughout the FTLD and MND spectrum that are not well delineated, especially among common forms of MND like PLS, ALS, and PMA. In this study, we aimed to determine the prevalence of ANXA11 proteinopathy in 379 autopsy cases, including FTLD-TDP and FTLD-MND and MND-TDP cases subclassified neuropathologically into MND forms of PLS, ALS, and PMA. Furthermore, we conducted genetic analyses to identify rare *ANXA11* variants in 159 cases, including all ANXA11-positive cases.

## Methods

### Case selection and neuropathologic assessment

Individuals with a neuropathologic diagnosis of FTLD-TDP, FTLD-MND, or MND-TDP (*N* = 379) were selected from the Mayo Clinic Florida brain bank for neurodegenerative diseases. For most autopsies, the brain was divided in the mid-sagittal plane, with the right hemibrain frozen for biochemical and genetic studies and the left hemibrain fixed in formaldehyde. Brain dissection followed a systematic sampling protocol for Alzheimer disease[62]. Coronal sections of the spinal cord were sampled at multiple levels. All brains underwent diagnostic evaluation by an experienced neuropathologist (D.W.D.). Paraffin-embedded 5-μm thick sections were mounted on glass slides and stained with hematoxylin and eosin (H&E). Neurofibrillary tangle and amyloid plaque pathologies were assessed using thioflavin S fluorescent microscopy. Lewy-related pathologies were assessed with α-synuclein immunohistochemistry (NACP, rabbit polyclonal, 1:3000, Mayo Clinic; or phospho-S129, EP1536Y, rabbit monoclonal, 1:40000, Abcam) and classified according to the McKeith criteria[61]. Immunohistochemistry for phospho-S409/410-TDP-43 (pTDP-43, mouse monoclonal, 1:5000, Cosmo Bio, Tokyo, Japan) was performed on sections of the middle frontal gyrus, primary motor cortex, posterior hippocampus (at the level of the lateral geniculate nucleus), basal forebrain, medulla, and spinal cord, when available. *C9orf72* repeat expansion status was evaluated with C9RANT immunohistochemistry. Through previous genetic studies and routine genetic characterization of *GRN* in FTLD-TDP A cases and *C9orf72* in all cases, pathogenic variants in known neurodegenerative disease genes were found in 113 of the 379 cases (62 *C9orf72*, 35 *GRN,* six *DCTN1*, two *CCNF*, two *TARDBP*, two *VCP*, two *TBK1*, one *OPTN/TBK1* and one *LRRK2*)[10, 18, 21, 24, 25, 27, 49, 73, 74, 92].

### Neuropathologic evaluation and subclassification of MND

Motor neuron loss was evaluated semiquantitatively on H&E-stained sections. Upper motor neuron loss was assessed in the motor cortex and lower motor neuron loss was assessed in the hypoglossal nucleus and anterior horn of the spinal cord, when available. MND was subclassified into three forms based on the relative extent of upper and lower motor neuron loss: moderate to severe upper motor neuron loss with no or mild lower motor neuron loss = primary lateral sclerosis (PLS-TDP/FTLD-PLS); moderate to severe upper and lower motor neuron loss, as well as cases with mild upper and lower motor neuronal loss = amyotrophic lateral sclerosis (ALS-TDP/FTLD-ALS); no or mild upper motor neuron loss with moderate to severe lower motor neuron loss = progressive muscular atrophy (PMA-TDP/FTLD-PMA).

Degeneration of the corticospinal tract (CST) was assessed on Luxol fast blue-periodic-acid Schiff’s stained sections for myelin changes and IBA-1 immunostained sections to assess microglial activation (rabbit polyclonal, 1:3000, Wako Chemicals, USA). The CST was evaluated at supratentorial levels, including motor cortex white matter and posterior limb of the internal capsule, and at infratentorial levels, including the middle cerebral peduncle in the midbrain, the medullary pyramid, and the lateral corticospinal tract (upon spinal cord availability). CST degeneration was observed at all available levels in cases with PLS and ALS but was often more prominent in infratentorial than supratentorial levels in cases with PMA.

### Annexin A11 immunostaining

All individuals were stained using immunohistochemistry for an anti-annexin A11 antibody (rabbit polyclonal, 1:1000, 10479-2-AP, Proteintech) in the basal forebrain, which included the corticomedial amygdala, basolateral amygdala, globus pallidus, and putamen. FTLD-MND and MND-TDP cases were also screened for ANXA11 in the motor cortex, medulla (hypoglossal nucleus), and anterior horn of the spinal cord (when available). The six most recently evaluated FTLD-TDP type C and all other ANXA11-positive cases were further screened and evaluated in the middle frontal gyrus, posterior hippocampus (including dentate gyrus, CA1/subiculum, parahippocampal gyrus, and inferior temporal cortex), and medulla (including inferior olivary nucleus and medullary tegmentum). Then, pTDP-43- and ANXA11-immunoreactive neuronal cytoplasmic inclusions and neurites were graded semiquantitatively (0 = absent, 1 = mild, 2 = moderate, 3 = marked, with 0.5 half steps when appropriate) blinded to case information (N.B.G.) in thirteen brain regions.

### Immunofluorescent staining

Immunofluorescent staining was performed using antibodies to pTDP-43 (mouse monoclonal, 1:200, Cosmo Bio) and ANXA11 (rabbit polyclonal, 1:200, 10479-2-AP, Proteintech). Deparaffinized sections were blocked with DAKO Protein Block plus Serum Free for 1.5 hours, then incubated with primary antibodies overnight at 4°C. After, sections were incubated with fluorescently labeled secondary antibodies (1:500, Alexa Fluor 488 and 647) for 1.5 hours at room temperature, then briefly counterstained with 1% Sudan Black. Representative images were obtained with a confocal microscope (LSM 880, Carl Zeiss).

### Clinical analyses

Clinical and demographic information was abstracted from patient records (N.B.G., S.F.R., K.A.J.). The following cognitive, behavioral, speech/language, psychiatric, pyramidal, and extrapyramidal features were recorded as present or absent: cognitive impairment, memory loss, behavioral disturbance, speech impairment (including dysarthria), aphasia, psychiatric disorder, pseudobulbar affect, spasticity, clonus, Babinski sign (extensor plantar reflexes), hyperreflexia, generalized weakness, LMN involvement demonstrated by electromyography (EMG), fasciculations (in the absence of significant sensory impairments and/or EMG-indicated peripheral neuropathy), muscle atrophy (affecting at least three muscles), vertical supranuclear gaze palsy, any other oculomotor disturbances, rigidity, myoclonus, limb apraxia, hypomimia, any parkinsonism, frequent falls, postural imbalance, and ataxia. Symptoms and signs not tested or recorded were excluded from analyses and not considered absent.

### Genetic analyses

We screened for nonsynonymous coding variants in all twelve exons of the *ANXA11* gene in 106 ANXA11-positive cases. First, *ANXA11* variants were queried from a previously generated whole-genome sequencing (WGS) dataset, which included 44 cases[74]. All fourteen coding exons of *ANXA11* were Sanger sequenced in the remaining 62 ANXA11-positive cases that were not included in the WGS dataset. Genomic DNA (n = 36) was prepared from cerebellum tissue using standard protocols with the Qiagen DNeasy Tissue kit. Polymerase chain reaction (PCR) was performed with the extracted DNA as template to amplify all fourteen exons of the *ANXA11* gene using primers designed to flanking intronic regions, tailed with M13 sequences. The PCR products were purified using AMPure magnetic beads (Agencourt Biosciences) on a Biomek FXP robotic workstation, then sequenced in both directions using M13 primers and the Big Dye Terminator v3.1 Cycle Sequencing kit (Applied Biosystems). Sequencing reactions were purified using CleanSEQ (Agencourt Biosciences) and run on an ABI3730xl Genetic Analyzer (Applied Biosystems). Sequencing analysis was performed using Sequencher 5.4.6 software (Gene Codes). All variants were confirmed in separate PCR and sequencing reactions. All variants were confirmed in separate PCR and sequencing reactions. For regions difficult to sequence due to a poly-T tract before and a GT-rich region after exon 6, cleaner reads were obtained by sequencing ANXA11 cDNA synthesized using oligo(dT)-primed reverse transcription from TRIzol/RNeasy-purified RNA kit from available brain tissue (n = 26). Three ∼700 bp amplicons were designed to provide complete coverage of the 1.5 kb coding sequence (505 codons) for the PCR and sequencing reactions mentioned above. To identify rare variants in *ANXA11*, variants with gnomAD-NFE allele frequency >0.001 were excluded.

### Statistical analyses

Mean and standard error of the mean were reported for all demographic data. Fisher’s exact tests and relative risk ± 95% confidence intervals were calculated using Koopman asymptotic scores to compare the prevalence of each clinical variable, as well as for genetic comparisons. Wilcoxon matched-pairs signed rank test with Holm step-down method for multiple comparisons were performed for within groups comparisons. Kruskal-Wallis one-way ANOVA with Dunn’s multiple pairwise comparisons post-hoc tests were performed for between groups comparisons. For frequentist tests, *P* < 0.05 was considered significant after multiple comparisons corrections. All frequentist tests and line plots were performed in GraphPad Prism 10. Hierarchical clustering analyses of pTDP-43 and ANXA11 inclusion data from thirteen brain regions were performed utilizing manhattan distance with ward.D2 linkage for row and column data. Average silhouette widths and Jaccard indices were calculated to validate cluster number and stability of clusters (Fig. S1a). These analyses were performed using the “cluster”, “ComplexHeatmap“, and “fpc” packages in R (version 4.4.3).

## Results

### Annexin A11 in FTLD-MND

Based on an immunohistochemical screening of ANXA11 in our autopsy cohort of 379 cases, ANXA11 inclusions were more frequently detected in FTLD-MND than in FTLD-TDP or MND-TDP (Table 1). Consistent with previous reports, ANXA11 inclusions were found in all FTLD-TDP type C cases and were rare in FTLD-TDP cases with other TDP-43 proteinopathies. Since ANXA11 inclusions were present in many FTLD-MND cases, we next determined if ANXA11 proteinopathy was associated with a specific form of MND by neuropathologically subclassifying MND-TDP into three major forms: PLS-TDP, ALS-TDP, and PMA-TDP. Unexpectedly, ANXA11 inclusions were present in most FTLD-PLS cases (Fig. 1; *n* = 38/40, 95%), uncommon in FTLD-ALS (*n* = 8/41, 20%) and rare in FTLD-PMA (*n* = 1/28, 4%). Further, while six ANXA11-positive FTLD-PLS cases had TDP type C, the remaining 32 cases had either TDP type B or an unclassifiable TDP-43 proteinopathy with abundant, pleomorphic NCIs and neurites characteristic of multiple TDP-43 subtypes, including TDP types A, B, C, and E.

**Figure 1.**
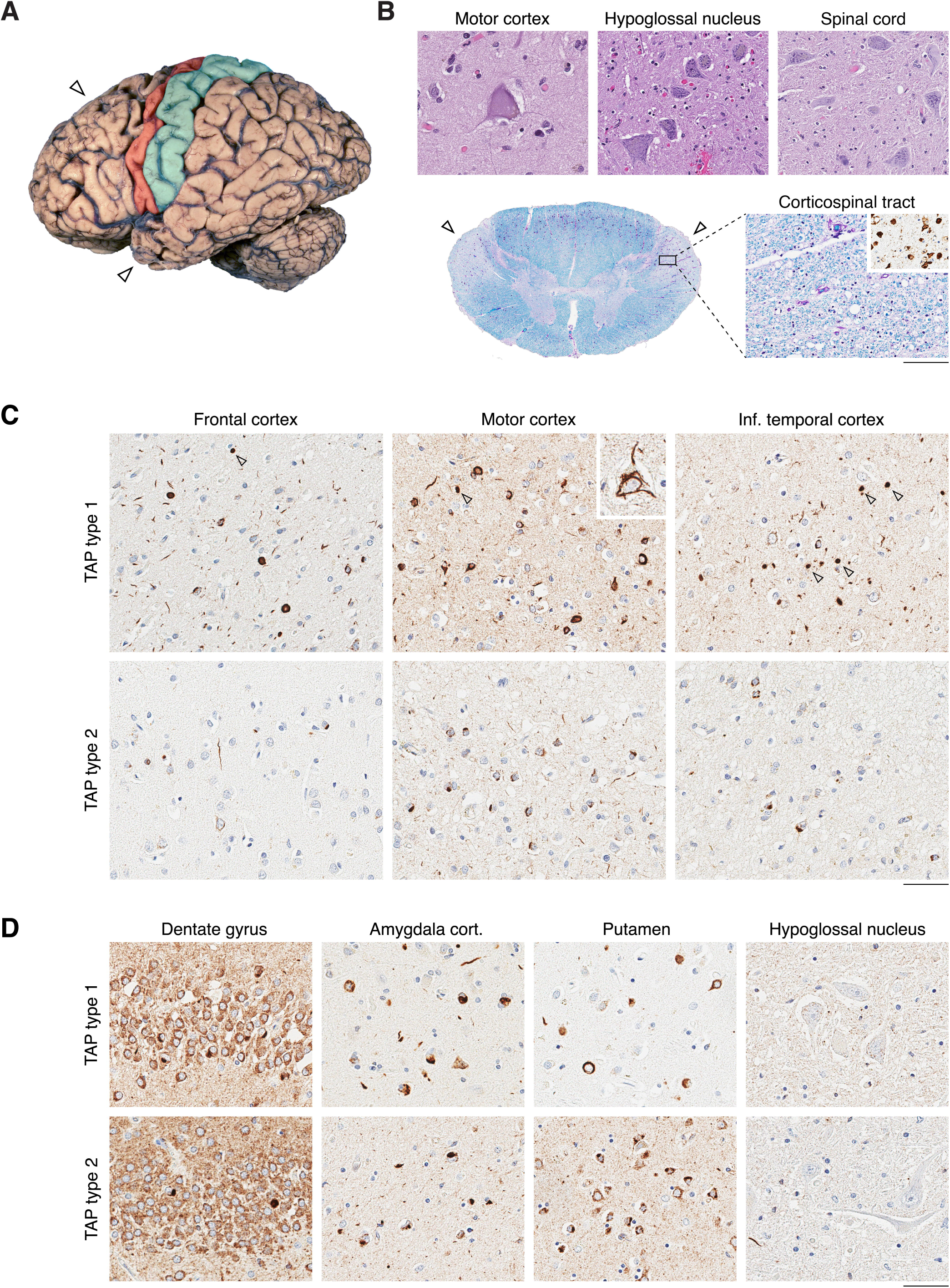
**FTLD-PLS pathology and ANXA11 in TAP types 1 and 2** (**A**) Macroscopic hallmarks of FTLD-PLS include severe frontal and anterior temporal lobe atrophy (hollow arrowheads) in addition to prominent primary motor cortex atrophy (highlighted in red). The primary sensory cortex is typically preserved (highlighted in cyan; P32). (**B**) Histologic hallmarks include degenerating Betz cells in primary motor cortex (P5), preserved neuronal populations in the hypoglossal nucleus (P3) and anterior horn cells of the spinal cord (P17), pale lateral corticospinal tracts (hollow arrowheads, P18) with Luxol fast blue-Periodic acid Schiff’s stain accompanied by microgliosis with IBA-1 immunohistochemistry (P18) in the thoracic spinal cord. (**C**) ANXA11 in neocortex of TAP types 1 and 2. Abundant pleomorphic neuronal cytoplasmic inclusions (NCI) and neurites in the frontal cortex (P3), motor cortex, and inferior temporal cortex (from P11) were indicative of TAP type 1. Small globular spheroids are also observed frequently in TAP type 1 neocortex (hollow arrowheads), as well as vermiform Betz cell NCIs (inset). Conversely, TAP type 2 features predominantly NCIs with sparse pleomorphic neurites in the frontal cortex, motor cortex, and inferior temporal cortex (from P18, P20, and 23, respectively). (**D**) ANXA11 in subcortical brain regions of TAP type 1 and 2. Pleomorphic dentate gyrus granule cell inclusions are more prominent in TAP type 1 than TAP type 2, and physiologic staining is abundant in most cases (from P11 and P27, respectively). NCIs and neurites are abundant in the corticomedial amygdala, wherein neurites appear thick and of variable length in TAP type 1, and fine or speck-like in TAP type 2 (from P3 and P32, respectively). TAP type 1 putaminal NCIs are often more diffusely abundant and appear ring-like (from P11), whereas NCIs are typically abundant in small groups and morphologically globular and/or granular (from P32). Lower motor neurons of the hypoglossal nucleus are usually spared in both types and sparse neurites can be observed in TAP type 2 (from P11 and P20, respectively). Scale bar(s) = 50um.

**Table 1.** Frequency of ANXA11-immunoreactive inclusions in FTLD-TDP, FTLD-MND, and MND-TDP (*N* = 379).

| Pathology | Subtype | N | # ANXA11 positive (%) | TDP-43 pathologic subtype |  |  |  |  |  | Unclassifiable |
| --- | --- | --- | --- | --- | --- | --- | --- | --- | --- | --- |
|  |  |  |  | A | B | C | D | E | P |  |
| FTLD-TDP |  | 174 | 57 (33%) | 13/107 | 2/14 | 42/42 | 0/2 | 0/1 | 0/6 | 0/2 |
| FTLD-MND |  | 109 | 47 (43%) |  |  |  |  |  |  |  |
|  | Progressive muscular atrophy (FTLD-PMA) | 28 | 1 (4%) | - | 0/23 | - | - | 0/3 | - | 1/2 |
|  | Amyotrophic lateral sclerosis (FTLD-ALS) | 41 | 8 (20%) | - | 6/33 | - | - | 0/4 | - | 2/3 |
|  | Primary lateral sclerosis (FTLD-PLS) | 40 | 38 (95%) | - | 15/17 | 6/6 | - | - | - | 17/17 |
| MND-TDP |  | 96 | 2 (2%) |  |  |  |  |  |  |  |
|  | Progressive muscular atrophy | 23 | 0 (0%) | - | 0/23 | - | - | - | - | - |
|  | Amyotrophic lateral sclerosis | 64 | 1 (1%) | - | 1/62 | - | - | 0/1 | - | 0/1 |
|  | Primary lateral sclerosis | 9 | 1 (11%) | - | 1/6 | - | - | - | - | 0/3 |

**Table 2.**
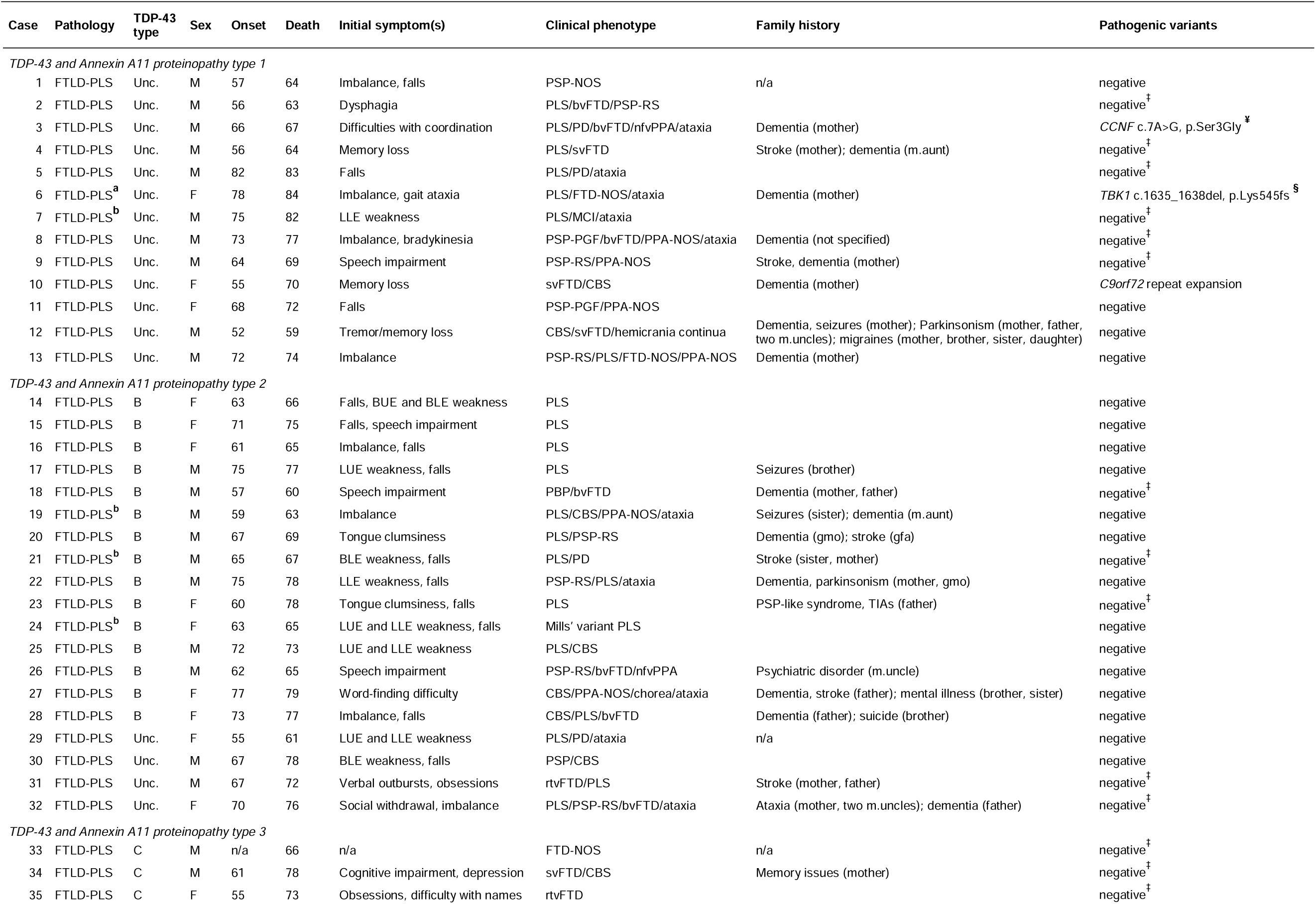
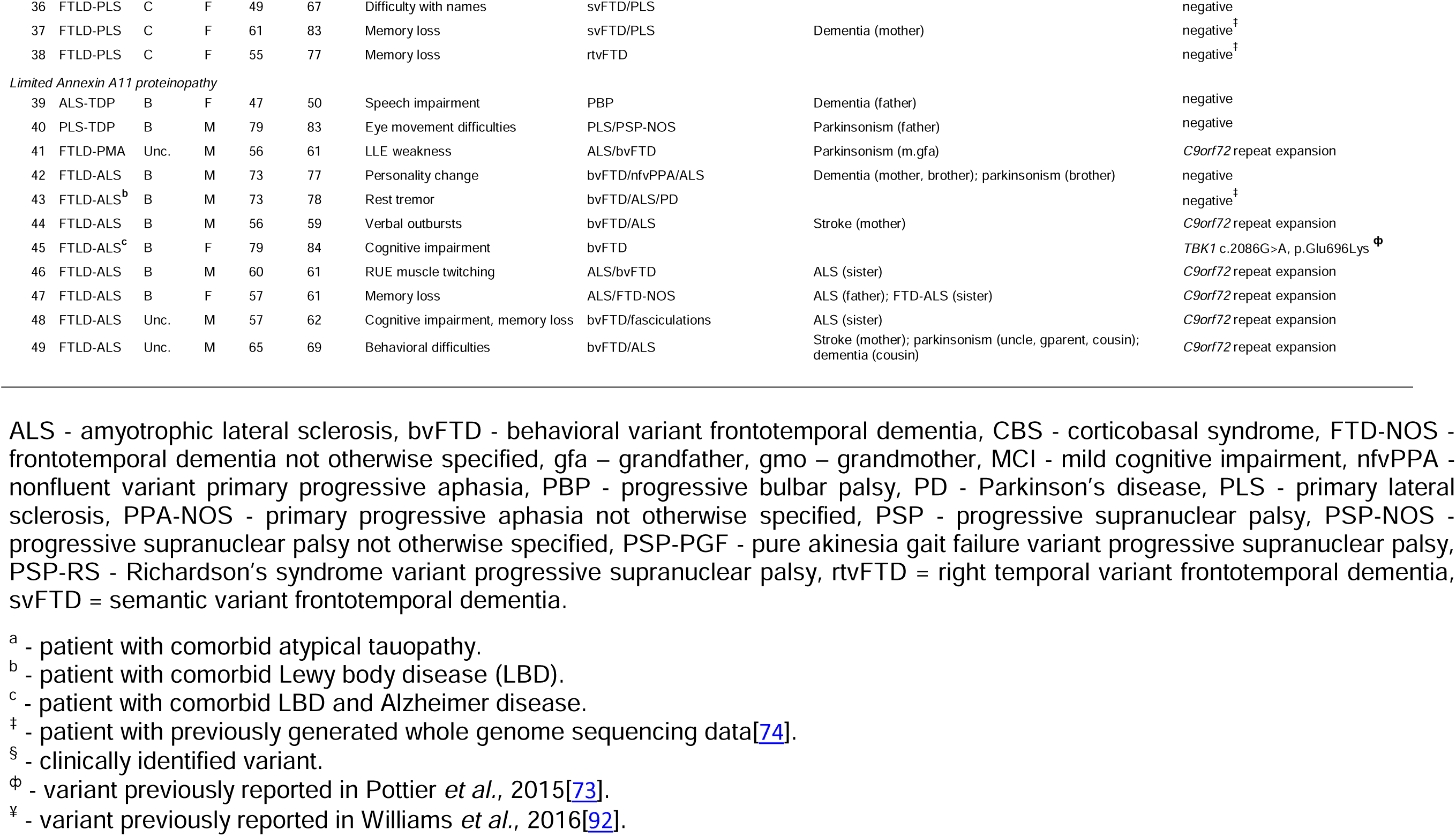
Clinical, pathologic, and genetic features of all ANXA11-positive FTLD-MND and MND-TDP cases.

### Pathologic subtypes of ANXA11 proteinopathy

Phospho-TDP-43 inclusions were relatively more abundant than ANXA11 inclusions in all ANXA11-positive cases with neuropathologic diagnoses of FTLD-PMA, FTLD-ALS, ALS-TDP, PLS-TDP, FTLD-TDP type A, and FTLD-TDP type B. Therefore, we provisionally classified the ANXA11 proteinopathy in these cases as limited ANXA11 proteinopathy (LAP). In contrast, pTDP-43 and ANXA11 inclusions were similarly abundant in all ANXA11-positive FTLD-PLS cases, as well as in all FTLD-TDP type C cases. Inclusions were consistently co-immunoreactive for pTDP-43 and ANXA11 in these cases with double immunofluorescent staining (Fig. 2). Given our observation of similarly abundant and co-immunoreactive pTDP-43 and ANXA11 inclusions in these cases, we provisionally classified the molecular pathology as a TDP-43 and ANXA11 proteinopathy (TAP). Then, in the 32 non TDP type C FTLD-PLS cases, we observed two distinct distributions and morphologies of inclusions. Therefore, we differentiated three subtypes of TAP based on distinct patterns in the abundance, distribution, and morphology of inclusions. Each TAP type had the following distinctive pathologic features with pTDP-43 and ANXA11 immunostains:

**Figure 2.**
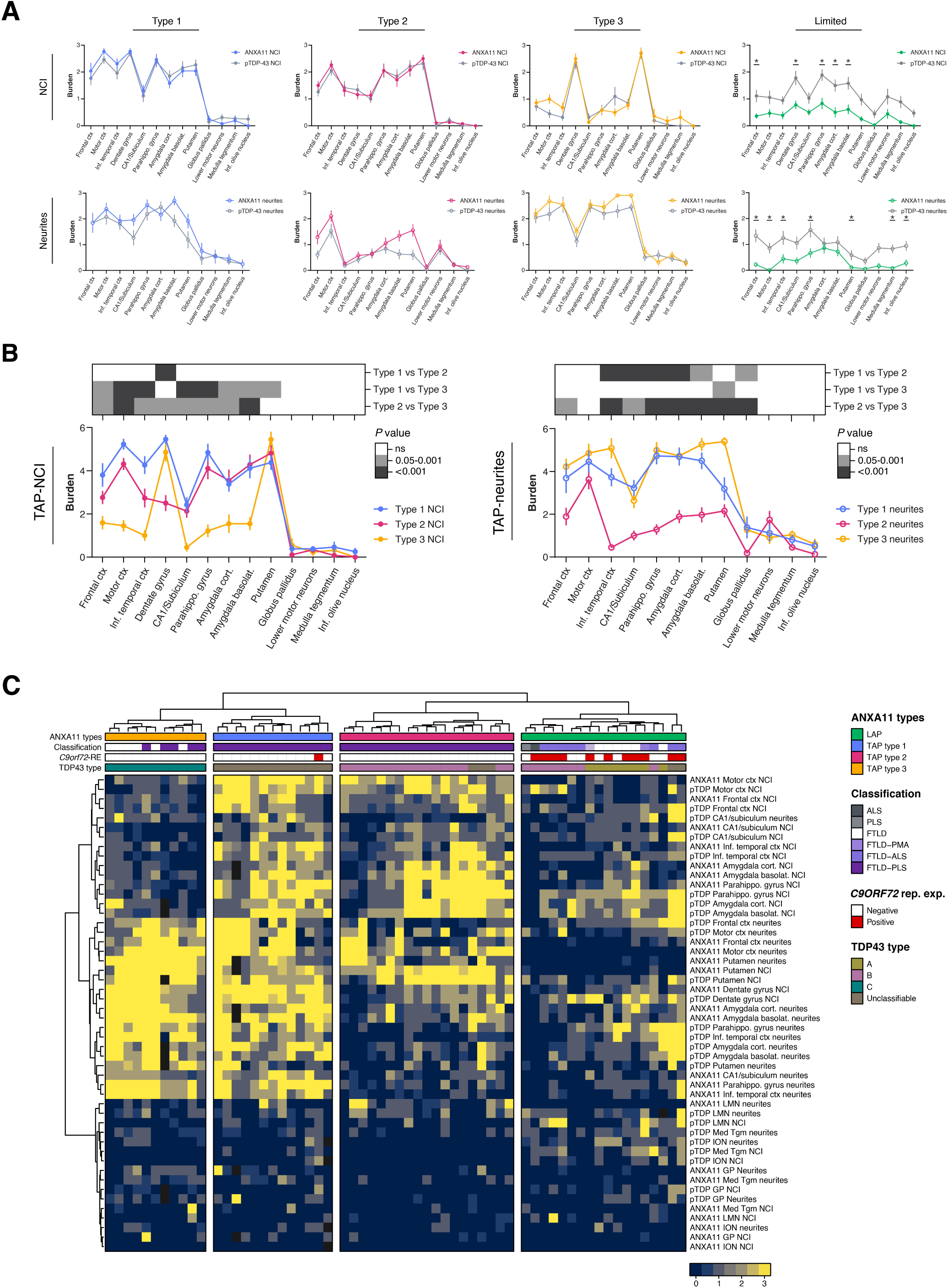
**pTDP-43 and ANXA11 immunofluorescence staining in TAP types 1-3** (**A**) Aggregates in the superficial layers of motor cortex and inferior temporal cortex show a high degree of co-immunoreactivity with pTDP-43 and ANXA11 antibodies across all three TAP types. Occasional ANXA11-immunoreactive neurites were not labeled with pTDP-43 in TAP type 1, and immunoreactivity varied with both antibodies in segments of some neurites in TAP type 3. Scale bar = 50um.

**TAP type 1** included thirteen FTLD-PLS cases (*n =* 13/40, 33%) with unclassifiable TDP-43 proteinopathy and was characterized by widespread and morphologically diverse neuronal cytoplasmic inclusions (NCIs) and neurites. NCIs were abundant in multiple neocortices and more numerous in superficial cortical layers. Perinuclear, ring-like aggregates were common in superficial NCIs as well as globular aggregates. On the other hand, deeper cortical NCIs were frequently small, round, and compact, and few were granulofilamentous. Skein- and comet-like NCIs were observed in Betz cells. Subcortical NCIs often appeared globular, granulofilamentous, and ring-like, and NCI morphologies were especially pleomorphic in basolateral amygdala. Four out of thirteen cases had neuronal intranuclear inclusions (NII) in neocortex or putamen with pTDP-43. NII were not observed with ANXA11. Glial cytoplasmic inclusions (GCI) were detected with pTDP-43 in the motor cortex white matter of four cases and in the medullary pyramid of just one of these cases. GCI were not clearly observed with ANXA11.

Neurites were abundant in multiple neocortical, limbic, and basal ganglia regions. Cortical neurites were observed in all cortical layers, though most abundant in superficial and middle, or pyramidal, cortical layers. Morphologically, cortical neurites typically ranged in appearance from short neurites of TDP type A to rope-like neurites of TDP type C and grain-like inclusions of TDP type E. Subcortical neurites were similarly pleomorphic and often appeared thicker than cortical neurites. Unusual globular spheroids, similar in size to nearby neurons, were regularly observed in superficial cortex and subcortical regions. Neurites were sparse in the globus pallidus, though few cases had abundant grain-like inclusions in globus pallidus fiber tracts.

**TAP type 2** included nineteen FTLD-PLS cases (*n =* 19/40, 48%), fifteen with TDP type B and four with an unclassifiable TDP-43 proteinopathy, and was characterized by many pleomorphic NCIs and limited neuritic pathology. Cortical NCIs were relatively more abundant in superficial cortical layers, though few cases had primarily deeper cortical NCIs. Most superficial NCIs appeared dense and round, and a subset had granular and compact features, while deeper cortical NCIs were predominantly granular. Unique speckled cytoplasmic inclusions were regularly observed in Betz cells and were variably labeled with pTDP-43 and ANXA11. A spectrum of granular and small, compact aggregates was also common among subcortical NCIs. NIIs were not observed with pTDP-43 or ANXA11. GCIs were detected with pTDP-43 in the motor cortex white matter of five out of nineteen cases and in the medullary pyramid of three of these cases. GCI were not clearly observed with ANXA11.

TAP type 2 neurites were most abundant in motor cortex and much less abundant in other neocortices. Motor cortex neurites often affected similar cortical layers as TAP type 1 neurites, though marginally less abundant. Motor cortex neurites were also less variable in appearance, ranging between those of TDP types A and C. Neurites in other neocortices were rare, but morphologically similar to neurites in motor cortex. Globular spheroids were rare and observed only in subcortical regions. Grain-like inclusions and thin neurites were prominent in limbic regions and the putamen, and rare in lower motor neuron regions.

**TAP type 3** included all 48 cases with TDP-43 type C and was characterized by a predominance of long rope-like neurites. NCIs were infrequent in most regions, except the dentate gyrus granule cells and putamen where they were highly abundant. Minimal NCIs were observed in lower motor neurons. Neurites were most prominent in superficial and middle cortical layers.

### Pathologic analyses

Next, we validated these ANXA11 proteinopathy subtypes by semiquantitatively scoring the abundance of pTDP-43 and ANXA11 NCIs and neurites across thirteen brain regions in cases with LAP and cases with TAP types 1, 2, and 3. Then, we compared the regional abundance of pTDP-43 and ANXA11 inclusions within each proteinopathy type (e.g. – we compared the abundance of pTDP-43 motor cortex NCIs to ANXA11 motor cortex NCIs within TAP type 1). Phospho-TDP-43 and ANXA11 NCIs or neurites were similarly abundant in all regions of TAP types 1, 2, and 3 (Fig. 2a). In LAP cases, ANXA11 NCIs and neurites were significantly less abundant than pTDP-43 NCIs and neurites in multiple regions.

Then, to understand the pathologic differences between TAP types, we summed each patient’s regional pTDP-43 and ANXA11 inclusion scores to generate regional TAP-NCI and TAP-neurite scores (Fig. 2b). For instance, to generate a patient’s motor cortex TAP-NCI score, we summed their motor cortex pTDP-43 NCI and motor cortex ANXA11 NCI scores. Intriguingly, while a similar abundance of TAP-NCIs was observed in most regions between TAP types 1 and 2, TAP-neurites were more frequent in TAP type 1 across numerous neocortical and subcortical regions (Fig. 2b). On the other hand, between TAP types 1 and 3, TAP-NCIs were significantly more abundant in TAP type 1 across many neocortical and limbic regions, and a similar abundance of TAP-neurites was observed in most regions. Between TAP types 2 and 3, neocortical and limbic TAP-NCIs were more abundant in TAP type 2; however, TAP type 3 had a greater abundance of TAP-NCIs in the dentate gyrus and TAP-neurites in the frontal cortex, inferior temporal cortex, limbic, and basal ganglia regions. Pertinently, motor cortex TAP-neurites were similarly abundant in all three TAP types, and motor cortex TAP-NCIs were more abundant in TAP types 1 and 2 compared to TAP type 3.

We then investigated whether our differentiation of four distinct types of ANXA11 proteinopathy could be validated by performing manhattan distance-based clustering of the semiquantitative ANXA11 and pTDP-43 inclusion data (Fig. 2C). Remarkably, our differentiation of four distinct types of ANXA11 proteinopathy was reproduced as each case was clustered into their previously designated ANXA11 proteinopathy type. ANXA11 proteinopathy types were poorly clustered using just the pTDP-43 inclusion data (data not shown).

NCIs and neurites in TAP types 1, 2, and 3 were co-immunoreactive for pTDP-43 and ANXA11 by double immunofluorescence staining of the motor cortex and inferior temporal cortex (Fig. 3). Occasional cortical neurites labeled more strongly for ANXA11 than pTDP-43 in TAP type 1 cases, and relatively prominent immunoreactivity for either protein was observed at the ends of long neurites in TAP type 3 cases.

**Figure 3.**
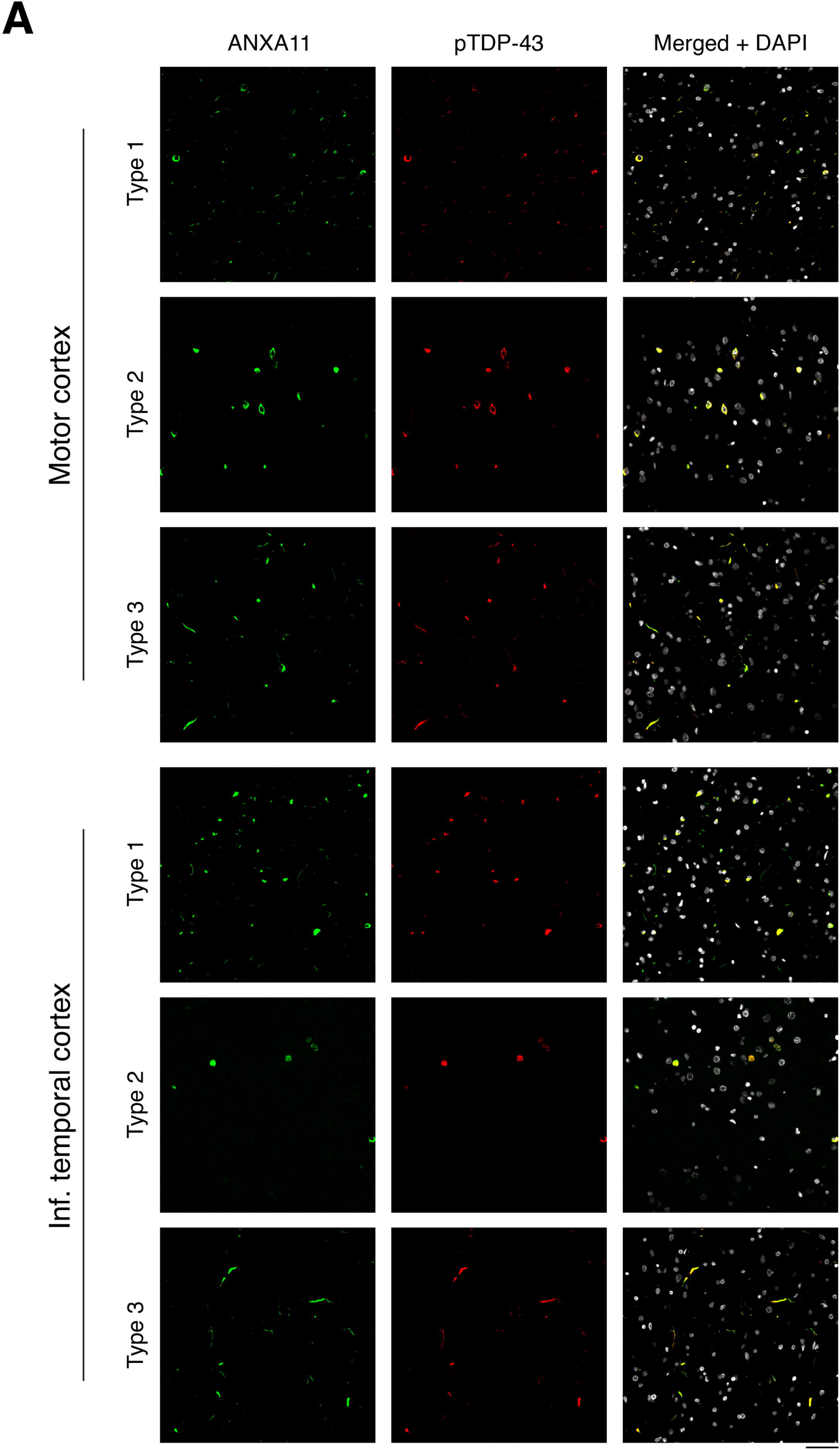
**Pathologic lesion analyses in ANXA11-related proteinopathy subtypes** (**A**) Line plots of the average burden of semiquantitatively scored NCIs and neurites observed with pTDP-43 and ANXA11 single immunostains in patients with TAP types 1, 2, 3 and limited ANXA11 proteinopathy (LAP; * *P* < 0.05). (**B**) Line plots of summed TDP-43 and ANXA11 NCI and neurite scores for each brain region and TAP type. Heatmap depicts *P*-values from pairwise comparisons between TAP types (Kruskal-Wallis one-way ANOVA with Dunn’s test for pairwise comparisons). (**C**) Heatmap visualization of semiquantitative pTDP-43 and ANXA11 lesion data hierarchically clustered into four groups using manhattan distance and ward.D2 linkage. Annotation layers (top) denote the ANXA11 proteinopathy subtype, pathologic classification, *C9orf72*-repeat expansion status (*C9orf72*-RE), and TDP-43 classification for each case.

### Clinical analyses

Given the additional context of ANXA11 proteinopathy, we investigated whether FTLD-PLS with TAP types 1 and 2 could be distinguished clinically (Fig. 4). Clinical summaries for the three ANXA11-negative FTLD-PLS cases are summarized (Supplementary information 1). Demographically, TAP type 1 commonly affected males (10/13 = 77%), while TAP type 2 affected similar proportions of males (10/19) and females (Supplementary table 1). The mean age of symptomatic onset (66 and 67 years-old) was similar between both types, but disease duration was marginally greater in TAP type 1 (6 years, *P=*0.0698) than in TAP type 2 (4 years, Table S2). Several TAP type 1 and 2 patients had a family history of dementia, parkinsonism, or stroke; however, none had a family history of MND.

**Figure 4.**
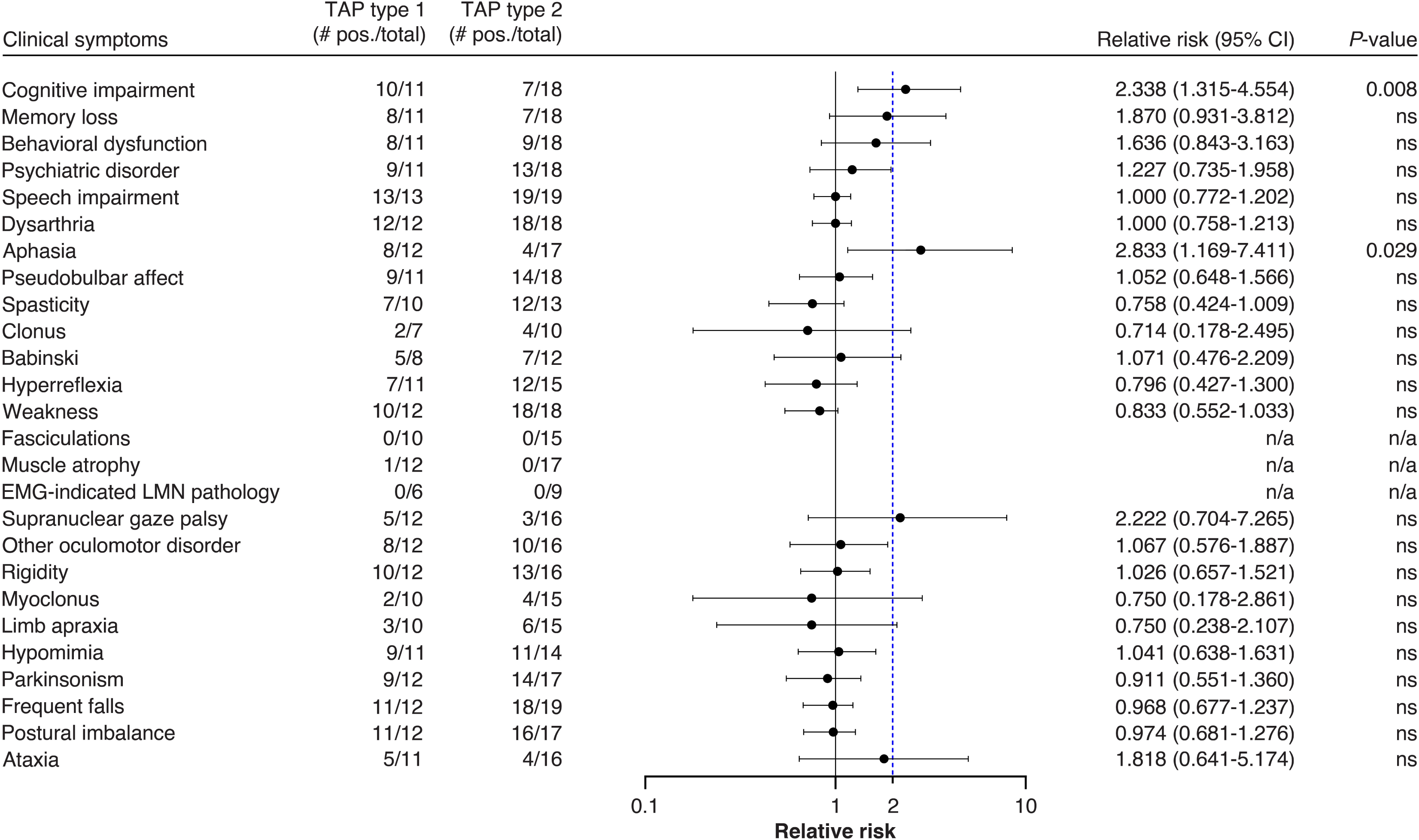
**Frequency and relative risk ratio of clinical features between TAP types 1 and 2** Relative risk (RR) ratio and 95% confidence intervals are plotted on logarithmic scale, blue dashed line denotes relative risk ratio of two. RR ratio was not calculated when symptom frequency was equal to zero.

At symptom onset, three TAP type 1 patients presented with memory loss, whereas no TAP type 2 patients reported memory loss at onset. Conversely, muscle weakness was noted in many TAP type 2 cases (8/18, *P*=0.050, RR=5.474) and in just one TAP type 1 case. Across both types, weakness often affected the lower extremities (8/9) or localized to the left hemibody (6/9) at symptom onset. Throughout the disease course, TAP type 1 had an increased relative-risk (RR) of developing cognitive impairment (10/11; *P*=0.008, RR=2.338) or aphasia (8/12; *P*=0.029, RR=2.833) than TAP type 2 patients (6/18, 4/17, respectively). Further, several TAP type 1 patients had svFTD, and one TAP type 2 patient had svFTD that primarily localized to the right temporal lobe, or right temporal variant FTD (rtvFTD). All TAP type 1 and 2 patients had speech disorders like dysarthria or anarthria.

Pertinently, UMN signs were common and LMN signs were rare in both TAP types. Additionally, no patients had atrophy or fasciculations in three or more muscles that could be solely attributed to LMN changes, and electromyographic (EMG) studies were negative for LMN involvement in all patients. On the other hand, extrapyramidal and other movement-related signs were prominent. One TAP type 1 patient received a DAT scan, which was abnormal, whereas three TAP type 2 patients received DAT scans and only one was abnormal. Neuroimaging was available in one TAP type 2 patient (P32) who underwent fluorodeoxyglucose-positron emission tomography (FDG-PET), which revealed left-predominant hypometabolism in frontal, motor, and anterior temporal cortices, and bilateral hypometabolism in the anterior corpus callosum, striatum, and posterior parietal cortices.

### Genetic analyses

Pathogenic *ANXA11* variants were not identified in any TAP case. Several rare *ANXA11* variants identified in LAP and ANXA11-negative cases were benign or likely benign (Supplementary material 1). Several variants in known neurodegenerative disease genes were identified in ANXA11-positive cases through routine screening. Within TAP type 1, one case (P6) carried a pathogenic *TBK1* variant (p.Lys545fs*), one case (P3) carried a *CCNF* variant (p.Ser3Gly), and one case (P10) carried a pathogenic *C9orf72* repeat expansion (*C9orf72*-RE)[92]. Within TAP type 2, no variant carriers were identified. Within TAP type 3, one FTLD-TDP case was identified with a *LRRK2* risk variant (p.Gly2019Ser)[18]. *C9orf72*-RE were frequently identified in cases with LAP (13/26), including many FTLD-MND (6/9) and MND-TDP patients (1/2). In addition, one case with FTLD-ALS and LAP carried a pathogenic *TBK1* variant (p.Glu696Lys), and another case with FTLD-TDP type A and LAP carried a *CCNF* variant (p.Arg392Thr)[73, 92]. Despite identifying ANXA11 proteinopathy in two pathogenic *TBK1* variant carriers, ANXA11 proteinopathy was not observed in a previously reported patient with pathogenic variants in both *TBK1* and *OPTN*[73]. Additionally, ANXA11 proteinopathy was not observed in carriers of pathogenic *GRN*, *DCTN1*, *TARDBP*, or *VCP* variants (*n =* 35, 6, 2, 2)[24, 25, 27, 75].

Intriguingly, *C9orf72*-RE were more common in FTLD-MND with LAP (6/9) than in ANXA11-negative FTLD-MND (12/109; *P*=0.0004; RR=6.056). On the other hand, *C9orf72*-RE were nominally more common in FTLD-TDP with LAP (6/15) than in ANXA11-negative FTLD-TDP type A (14/94; *P*=0.0306; RR=2.686), but not when compared to ANXA11-negative FTLD-TDP type B (2/12; *P*=0.2357; RR=2.400).

## Discussion

In this study, we identified ANXA11 proteinopathy in over 40% of FTLD-MND and in 95% of FTLD-PLS cases, of which the majority had TDP type B or an unclassifiable TDP-43 proteinopathy. We described two novel patterns of ANXA11 proteinopathy in FTLD-PLS and confirmed previous reports of ANXA11 proteinopathy in all TDP type C cases. Further, we did not identify pathogenic *ANXA11* variants in any ANXA11-positive case. These findings contrast with previous reports, where ANXA11 proteinopathy has primarily been observed in TDP type C and in carriers of pathogenic *ANXA11* variants[7, 33, 47, 67, 78, 79, 82, 87, 93]. Thus, we demonstrated novel forms of ANXA11 proteinopathy strongly associated with FTLD-PLS, but not with TDP type C or pathogenic *ANXA11* variants.

Recent studies demonstrated that TDP-43 and ANXA11 are similarly abundant and colocalized in inclusions of TDP type C, a disease associated clinically with svFTD[78, 93]. Using cryo-EM, researchers also identified that TDP-43 and ANXA11 co-assemble into heteromeric amyloid filaments in TDP type C. These findings suggested that the molecular pathology of TDP type C was a TDP-43 and ANXA11 proteinopathy[7]. In the present study, pTDP-43 and ANXA11 were similarly abundant and consistently co-immunoreactive in inclusions of all ANXA11-positive FTLD-PLS cases. Therefore, we propose that the molecular pathology underlying these cases is a TDP-43 and ANXA11 proteinopathy (TAP) and subclassify three types of TAP based on pathologic differences. Pathologically, TAP type 1 consists of pleomorphic NCIs and neurites and comprises most FTLD-PLS cases with unclassifiable TDP-43 proteinopathy. TAP type 2 is an NCI-predominant pathology and comprises most FTLD-PLS cases with TDP type B and few with unclassifiable TDP-43 proteinopathy. Lastly, TAP type 3 is a neurite-predominant pathology and includes all cases with TDP type C. Clinically, TAP type 1 patients are frequently male and more often have cognitive impairment or aphasia than TAP type 2 patients. Moreover, TAP type 2 patients more often present initially with muscle weakness affecting the left leg or hemibody. Intriguingly, several patients with TAP types 1, 2, and 3 presented with overlapping clinical features of svFTD and PLS. Therefore, we believe our delineation of novel TDP-43 and ANXA11 proteinopathies with shared, yet distinct, clinical and pathomolecular characteristics establishes a link between FTLD and PLS within the broader FTLD-MND spectrum.

Autopsy studies of FTLD-PLS are scarce. However, it is conceivable that others may have described pathology that retrospectively resembles TAP types 1 or 2. Several case reports and case series have described FTLD-PLS with TDP type B, TDP type C, and unclassifiable TDP-43 proteinopathies resembling TDP type A[4, 31, 42, 60, 93, 95]. One study described two ‘PLS with FTLD’ cases with difficult to classify TDP-43 proteinopathy that bare strong resemblance to our TAP type 1[42]. Examination of ANXA11 in other autopsy cohorts is required to validate our findings and better understand the prevalence of TAP types 1 and 2.

Given that we did not identify pathogenic *ANXA11* variants in any TAP case, it is worth considering how proteinopathies reported in pathogenic *ANXA11* variant carriers may be distinguished from sporadic TAP. Abundant TDP-43 and ANXA11 inclusions have also been described in carriers of pathogenic *ANXA11* variants; however, these studies have not consistently described TDP-43 and ANXA11 co-immunoreactivity in inclusions and have also demonstrated separate aggregates of each protein in individual or co-labeled inclusions[33, 47, 59, 67, 78, 79, 87]. Hence, we identified widespread TDP-43 and ANXA11 co-immunoreactivity in inclusions as a characteristic feature of TAP. Though UMN degeneration has been described in *ANXA11* variant carriers, most also have significant LMN degeneration[78, 79, 82, 87]. Further, several studies have also reported inclusion body myopathy (IBM) with differing degrees of UMN and LMN signs[33, 47, 59, 67]. Therefore, in the same way that *MAPT* variant-associated tauopathies may share features with sporadic tauopathies, *ANXA11* variant-associated proteinopathies may also share features with sporadic TAP, like UMN degeneration[1, 8, 15, 17, 43, 48, 84]. However, given that TAP is considerably more common than pathogenic *ANXA11* variants, pathologic interpretation of *ANXA11* variants requires consideration for whether the patient has TAP. Indeed, one p.G38R variant carrier, who had FTLD with UMN-predominant degeneration and TDP-43 and ANXA11 inclusions resembling TDP type A, may have had TAP type 1 instead[87]. While reports of *ANXA11* variants in clinical FTD cohorts suggest that the clinicogenetic spectrum of *ANXA11* variants might extend beyond FTLD-MND, MND-TDP, or IBM, autopsy studies are lacking in these patients[39, 46, 49, 66, 83, 96]. Therefore, more autopsy studies of *ANXA11* variant carriers are required to better understand these differences.

Based on our findings, it is unlikely that genetic variants in *ANXA11* explain the occurrence of ANXA11 proteinopathy in our autopsy cohort. We did identify pathogenic *TBK1* variants in a TAP type 1 patient and an FTLD-ALS patient with limited ANXA11 proteinopathy (LAP). A previous study also reported a pathogenic *TBK1* variant carrier with FTLD-ALS and sparse ANXA11 proteinopathy[78]. That being said, one patient with pathogenic variants in both *OPTN* and *TBK1* did not have ANXA11 proteinopathy, suggesting that ANXA11 proteinopathy might not be specific for pathogenic variants in these particular genes[73]. The two *CCNF* variant carriers included in our study were also both ANXA11-positive (one TAP type 1 and one LAP); however, future studies in additional mutation carriers are needed to determine whether ANXA11 proteinopathy is a consistent feature of *CCNF*-related disease. Notably, pathogenic repeat expansions in *C9orf72* (*C9orf72*-RE) were detected in many LAP patients, as well as one TAP type 1 patient. In particular, *C9orf72*-RE were more common in FTLD-MND with LAP than in ANXA11-negative FTLD-MND. Consistent with our findings, *C9orf72*-RE were recently described in two FTLD-ALS patients with sparse ANXA11 proteinopathy as well[78]. Hence, these findings could suggest that the expansion might be present in certain patients with ANXA11 proteinopathy, these initial observations warrant independent validation and further investigation. Nevertheless, despite our detection of several pathogenic variants (e.g., *TBK1*, *CCNF*, and *C9orf72*) in patients with TAP type 1 and LAP, most cases with ANXA11 proteinopathy remain genetically unexplained.

Proteinopathies besides TDP-43 or ANXA11 have been associated with FTLD-PLS, including tau proteinopathies, like PSP, CBD, and globular glial tauopathy (GGT), and neuronal intermediate filament inclusion disease with FET proteinopathy (NIFID-FET)[2, 9, 11, 12, 35, 44, 53, 63–65, 88]. FTLD-PLS patients with tau or FET proteinopathy present with features of FTD, atypical parkinsonism, and PLS, like many patients with TAP types 1 and 2. Therefore, future efforts to clinically differentiate FTLD-PLS with different proteinopathies are necessary to better identify patients with TAP types 1 and 2 clinically. Additionally, our identification of TAP proposes a distinct molecular signature for future biomarker discovery studies to explore in hopes of clinically detecting cases with different molecular pathologies, including cases with only TDP-43 proteinopathy.

Our findings may improve recognition of the clinicopathologic overlap of FTD and PLS. At present, FTD is not listed in several clinical PLS criteria, and recent consensus diagnostic criteria for PLS state that only a minority of PLS cases have widespread brain involvement[29, 76, 89]. Despite this, many studies have reported the clinical overlap of FTD and PLS, and nearly 80% of pathologic PLS cases had FTLD in our study[19, 20, 22, 23, 28, 31, 32, 34, 42, 72, 85, 93, 95]. Therefore, clinical and neuropathologic perspectives appear to be at odds and may reflect different criteria used to define PLS in both settings. FTD and widespread neurodegeneration may be more frequent in pathologic PLS than previously appreciated clinically, and revisions to clinical PLS criteria may be necessary to reflect the underlying pathology and to better detect FTLD-PLS cases with TAP.

Additionally, overlap in vulnerable brain regions across TAP types may indicate an overlap in underlying pathomechanisms. Physiologically, ANXA11 plays a prominent role in axonal transport of membrane-bound and membraneless organelles[50–52, 67, 81, 83]. Along with this, pathology was widespread and not associated with significant LMN degeneration in all TAP types. Therefore, it is possible that specific projections from UMNs, which reside in agranular motor cortex, may be vulnerable to TAP. Conventional hypotheses proposed that motor neuron loss in ALS follows a dying-back pattern, wherein corticospinal tracts of UMNs are affected secondarily to loss of spinal α-motor neurons[13]. However, corticospinal projections are heterogeneous and estimates suggest that only as many as 20% terminate directly on spinal α-motor neurons and that many also terminate on spinal interneurons, medium-sized putaminal neurons, and neurons of other neocortices[58]. Following this, putaminal NCIs were consistently abundant in all three TAP types, and pathology was observed in several non-motor neocortices in FTLD-PLS cases, especially in those with TAP type 1. Therefore, our findings in TAP cases with FTLD-PLS suggest that projections from UMNs to neurons in the putamen and other neocortices may be more vulnerable than projections to spinal α-motor neurons and interneurons. The abundance of motor cortex neurites in TAP may bolster this idea; though, conversely, the paucity of clinical and pathologic PLS in TAP type 3 may suggest that other brain regions are vulnerable. Therefore, spatial proteomic and transcriptomic investigations of the brain, as well as animal and cellular models of TDP-43 and ANXA11 dysfunction, may be required to clarify our understanding of selectively vulnerable neuronal populations and the directionality of pathological spread in ANXA11 proteinopathies.

It has long been debated whether PLS is simply a *forme fruste* of ALS or a unique disease entity since both diseases affect UMNs and their descending corticospinal tracts. Here, we propose that the molecular pathology in the majority of PLS is distinct from ALS and consists of a TDP-43 and ANXA11 proteinopathy. We acknowledge that our proposed classification of TAP as a distinct molecular pathology requires validation. While it has been shown that TDP-43 and ANXA11 form heteromeric amyloid filaments in TAP type 3, the structure of amyloid filaments in TAP types 1 and 2 is not known[7]. Future efforts to elucidate these filament structures may provide additional clarity into ANXA11-related disorders and the molecular pathology of the FTLD-MND disease spectrum.

Our study includes several important limitations. First, our autopsy cohort recruits many patients from tertiary medical centers who present with atypical parkinsonian disorders, like progressive supranuclear palsy (PSP) and corticobasal syndrome (CBS), which were common clinical diagnoses in TAP types 1 and 2 patients. Therefore, our finding of different ANXA11 proteinopathies may, in part, be due to different referral biases in our autopsy cohort compared to previously examined cohorts[78]. Second, patient records were not collected in a systematic, prospective manner and clinical diagnoses were made based on a retrospective review of patient records. Third, despite the paucity of lower motor neuron signs and pathology in FTLD-PLS associated with TAP, EMG testing was not obtained on every FTLD-PLS patient in this study. Nevertheless, this study provides a comprehensive examination of ANXA11 across the spectrum of FTLD and MND, including several major forms of MND pathology, and thereby contributes significant new insights into the role of ANXA11 in disease and the underlying pathology of FTLD-PLS.

## Conclusion

The present study revealed that ANXA11 proteinopathy is present in the majority of patients with PLS pathology and established several distinct clinical and pathologic subtypes of a novel molecular pathologic entity, called TDP-43 and ANXA11 proteinopathy (TAP). As PLS was conventionally difficult to identify clinically, these findings suggest that diagnostic and therapeutic strategies targeting ANXA11 may lead to important mechanisms of detection and treatment for patients with PLS.

## Acknowledgments

We express our sincere gratitude to the patients and their families for their agreement to brain donation. We would like to acknowledge the continuous commitment, technical support, and teamwork offered by Monica Castanedes-Casey, Whitney I. Davis, Nathan S. Perez, Virginia R. Phillips, and Linda G. Rousseau. This work was supported by the Mayo Clinic Alzheimer’s disease Research Center (P30 AG062677), R01-AG037491, UG3 NS103870, Fund for Scientific Research Flanders (FWO) grant # G064223N, CurePSP Foundation, and support from the Mayo Foundation.

## Ethics declarations

### Conflict of interest

KAJ and RR are associated editors at Acta Neuropathologica. RR receives invention royalties from a patent related to progranulin. KAJ is an Associate Editor of Annals of Clinical and Translational Neurology. The remaining authors declare no competing interests.

### Ethical approval and consent to participate

This study was approved by the Mayo Clinic and University of Antwerp Institutional Review Boards. All procedures performed in studies involving human participants were in accordance with the ethical standards of the institutional and/or national research committee and with the 1964 Helsinki Declaration and its later amendments or comparable ethical standards. All autopsies were obtained with consent from the legal next-of-kin or someone legally authorized to make this decision and conducted with the explicit assumption that the tissue will be used for diagnostic evaluation and research. The collection of tissue samples was approved by the Mayo Clinic Institutional Review Board.

