## Supplementary information 1 for "Expanding the spectrum of annexin A11 proteinopathy in frontotemporal lobar degeneration and motor neuron disease"

Supplementary Material

Supplementary Information 1 Clinical features of two ANXA11-negative FTLD-PLS cases.

**Case 1**—This right-handed female developed vision impairment and eyelid droop at age 74. Thereafter, she developed prominent speech impairment and increased emotionality, including inappropriate crying. A neurologic assessment at age 76 showed intact cognition with severe dysarthria, sialorrhea, vertical more than horizontal gaze limitation, marked nuchal rigidity with moderate postural hyperextension, and impaired balance on pull test. Strength, muscle bulk, and deep tendon reflexes were normal and there were no fasciculations. Performance on the Unified Parkinson’s disease rating scale (UPDRS) showed mild impairment, with a motor score of 15. At age 77, her ability to open her mouth reduced and she developed dystonic spasms and increased tone in the masseter and temporalis muscles bilaterally. She also developed walking impairments at this age. Over the next year, her cognition declined, she began to lose weight, and she developed significant swallowing difficulties. Her vertical gaze also worsened and a neurologic assessment was notable for severe anterocollis and increased fatigue. Repeat UPDRS testing showed little progression, with a motor score of 18. She died at the age of 78 with a clinical diagnosis of PSP.

**Case 2**—This right-handed female presented at age 62 with word-finding and vision difficulties. A neurologic assessment revealed mild difficulty with verbal expression, marked abnormalities of visual tracking and fixation, diffuse hyperreflexia, and imbalance. A neuroophthalmologic evaluation revealed extraocular movement abnormalities affecting pursuits and saccades suggestive of a disorder affecting the frontal eye fields without ophthalmoparesis. She was also found to have glaucoma. At age 63, she developed changes in her speech and cognitive processing. Her speech was halting, soft, monotone, and slurred. A neurologic assessment was remarkable for numerous neurologic alterations, including reduced verbal and semantic fluency, short-term memory deficits, personality change, left greater than right-sided bradykinesia, axial paratonia, and limb ataxia. Hoffmann’s sign and crossed adductor reflexes were present, but toes were downgoing. She also had mild right facial weakness in a lower motor neuron nerve distribution with synkinetic movements when blinking or smiling, and right hemifacial spasm with platysma involvement. At age 64, she developed intermittent episodes of uncontrollable laughing and crying, and dysarthria. On the Montreal cognitive assessment (MoCA), she scored 22/30, losing points on executive/visuospatial, naming, attention, language, delayed recall, and orientation domains. She began experiencing right arm and left leg weakness. On repeat MOCA testing, she scored 19/30, performing worse on language and executive/visuospatial domains. She also admitted to suicidal ideations and severe depression. She had a family history of ALS in her father and her sister reported difficulties using her legs. She died at the age of 64 with a clinical diagnosis of PSP and PPA-not otherwise specified.

Supplementary table 1 Demographics of TAP types 1 and 2

| **Feature** | **TAP type 1 (*n* = 13)** | **TAP type 2 (*n* = 19)** | ***P-*value** |
| --- | --- | --- | --- |
| # male (%) | 10 (77%) | 10 (53%) | ns |
| Age at onset (yrs, mean±SEM) | 66 ± 3 | 67 ± 2 | ns |
| Disease duration (yrs, mean±SEM) | 6 ± 1 | 4 ± 1 | 0.0698 |
| Braak stage (median±IQR) | 2 (1,2) | 2 (2,3) | ns |
| Thal phase (median±IQR) | 0 (0,II) | I (0,II) | ns |

Supplementary table 2 Cases screened for variants in *ANXA11*

| **Disease** | **Total # of cases** | **# screened for *ANXA11* variants** |
| --- | --- | --- |
| *ANXA11-positive cases* |  |  |
| TAP type 1 | 13 | 13/13 |
| TAP type 2 | 19 | 19/19 |
| TAP type 3 | 48 | 48/48 |
| Limited ANXA11 proteinopathy | 26 | 26/26 |
| *ANXA11-negative cases* |  |  |
| FTLD-TDP type A | 94 | 29/94 |
| FTLD-TDP type B | 12 | 0/12 |
| FTLD-TDP type D | 2 | 0/2 |
| FTLD-TDP type E | 1 | 1/1 |
| FTLD-TDP type P | 6 | 0/6 |
| FTLD-TDP type unclassifiable | 2 | 0/2 |
| FTLD-PMA type B | 23 | 7/23 |
| FTLD-PMA type E | 3 | 2/3 |
| FTLD-PMA type unclassifiable | 1 | 1/1 |
| FTLD-ALS type B | 28 | 9/28 |
| FTLD-ALS type E | 4 | 2/4 |
| FTLD-ALS type unclassifiable | 1 | 0/1 |
| FTLD-PLS type B | 2 | 2/2 |
| PMA-TDP type B | 23 | 0/23 |
| ALS-TDP type B | 61 | 0/61 |
| ALS-TDP type E | 1 | 0/1 |
| ALS-TDP type unclassifiable | 1 | 0/1 |
| PLS-TDP type B | 5 | 0/5 |
| PLS-TDP type unclassifiable | 1 | 0/1 |
